# Chondroitin sulfate, dermatan sulfate, and hyaluronic acid differentially modify the biophysical properties of collagen-based hydrogels

**DOI:** 10.1101/2023.05.22.541626

**Authors:** Marcos Cortes-Medina, Andrew R. Bushman, Peter E. Beshay, Jonathan J. Adorno, Miles M. Menyhert, Riley M. Hildebrand, Shashwat S. Agarwal, Alex Avendano, Jonathan W. Song

## Abstract

Fibrillar collagens and glycosaminoglycans (GAGs) are structural biomolecules that are natively abundant to the extracellular matrix (ECM). Prior studies have quantified the effects of GAGs on the bulk mechanical properties of the ECM. However, there remains a lack of experimental studies on how GAGs alter other biophysical properties of the ECM, including ones that operate at the length scales of individual cells such as mass transport efficiency and matrix microstructure. Here we characterized and decoupled the effects of the GAG molecules chondroitin sulfate (CS) dermatan sulfate (DS) and hyaluronic acid (HA) on the stiffness (indentation modulus), transport (hydraulic permeability), and matrix microarchitecture (pore size and fiber radius) properties of collagen-based hydrogels. We complement these biophysical measurements of collagen hydrogels with turbidity assays to profile collagen aggregate formation. Here we show that CS, DS, and HA differentially regulate the biophysical properties of hydrogels due to their alterations to the kinetics of collagen self-assembly. In addition to providing information on how GAGs play significant roles in defining key physical properties of the ECM, this work shows new ways in which stiffness measurements, microscopy, microfluidics, and turbidity kinetics can be used complementary to reveal details of collagen self-assembly and structure.

## 1. Introduction

The extracellular matrix (ECM) is vital to the maintenance of tissue as it confers cell signaling cues and is a semiporous barrier to solute and fluid transport.^1^ Fibrillar Type I collagen is the most abundant protein in the ECM. Thus, collagen is widely used by tissue engineers, materials scientists, and biologists to recapitulate *in vitro* the physical features of native *in vivo* tissue.^2^ Cells interact locally with collagen via specialized cell surface receptors such as integrins.^3^ Collagen can also be degraded by cell-secreted matrix metalloproteinases to help orchestrate tissue morphogenesis and organization.^4^

In addition to collagen, glycosaminoglycans (GAGs) are integral to the structure of the ECM and connective tissue.^5,6^ Among the GAG types, hyaluronic acid (HA), chondroitin sulfate (CS), and dermatan sulfate (DS) are the most prevalent in the body. These molecules are upregulated in pathologies characterized by an overabundance of fibrous connective tissue or desmoplasia such as cancer and fibrosis.^7-10^ HA is the only non-sulfated GAG and has a propensity to imbibe fluid and swell.^11,12^ Its role in disease pathogenesis and progression includes increased mechanical stress accumulation and physical barrier to molecular therapeutics.^13,14^ HA also alters cell signaling, either directly through its receptors CD44 and RHAMM, or indirectly by immobilizing growth factors and chemokines to generate concentration gradients from the ECM.^15^ Unlike HA, CS and DS are found predominantly as chains covalently linked to core proteins to form proteoglycans.^16,17^ CS and DS have similar structures except for the D-glucuronic acid units being changed to L-iduronic acid in DS.^18^ Like HA, the presence of CS and DS in the ECM alters tissue swelling, fluid retention, cell signaling.^16,19,20^

Collagen fiber assembly is a forward forming entropic reaction that is controlled by pH and temperature.^21^ Most of the physical and structural properties of collagen hydrogels are controlled by the architecture formed by the mesh of fibers.^22,23^ Yet, cells sense and respond to cues from multiple constituents within the ECM. Therefore, GAGs have been used to alter the bioactivity of collagen hydrogels. Moreover, GAGs have been used to modify the collagen polymerization kinetics and resultant fiber properties of composite hydrogels.^2,24,25^ Since HA, CS, and DS have some similar characteristics, one motivating factor is to understand if the presence of these GAG molecules exert distinct changes to the physical and structural properties of collagen.

Most prior experiments studying the mechanics of composite gels of GAGs and collagen have focused on their effects on bulk mechanical properties such as tissue-level stiffness. However, cells respond to fiber level features such radius, density, alignment, and matrix pore size. These physical properties of the ECM have been shown to mediate cell adhesion, clustering, and cancer cell invasion.^26-30^ Recently, our group showed that HA enhances interstitial flow-mediated angiogenic sprouting through its alterations to collagen ECM stiffness and pore size in a microfluidic-based tissue analogue model.^31^ This result demonstrates the importance of ECM composition and physical properties in controlling angiogenesis in the presence of biomolecular transport and fluid mechanical stimulation due to interstitial flow. Collectively, these examples emphasize the importance of investigating the collective effects of multiple ECM parameters on cell responses, rather than one aspect in isolation, such as ECM composition or stiffness.

Here we investigate and compare key mechanical, transport, and microstructural properties of collagen hydrogels laden with either CS, DS, or HA. We implement an integrated experimental biophysical characterization scheme for hydrogels previously developed by our group that profiles and decouples multiple ECM physical properties, including stiffness, hydraulic permeability, and matrix microarchitecture, using mechanical indentation testing, microfluidics, and confocal reflectance imaging and analysis respectively.^24^ As expected, HA significantly increased the indentation modulus of collagen hydrogels while surprisingly CS and DS had no effect. Addition of CS, DS, and HA decreased hydraulic permeability of collagen-based hydrogels. Interestingly, DS and HA significantly increased both collagen matrix pore size and fiber radius while CS had no effect on fiber radius and significantly decreased pore size. Lastly, we characterized collagen aggregate formation with turbidity measurements, which revealed that CS, DS, and HA differentially modify collagen polymerization kinetics. Collectively, this study details an enhanced understanding of the behavior of collagen assembly in the presence of different GAG constituents, which is relevant for understanding *in vivo* assembly of ECM.^22,32^ This work also clarifies how supplementation of collagen with different GAGs leads to the development of hydrogel systems of distinct structural and mechanical properties. This outcome enables improved design and utilization of collagen-based scaffolds used for tissue engineering of tailored composition, mechanical properties, molecular availability due to mass transport, and microarchitecture.

## 2. Materials and Methods

### 2.1. Preparation and Casting of Collagen Hydrogels

The collagen hydrogels used in the present study were prepared following the manufacturers ‘ instructions. In brief, rat tail type I collagen stored in acidic solution (Corning Life Sciences) was neutralized to a pH of 7.4 using sodium hydroxide and a 10X phosphate-buffered saline solution. The final concentration of the collagen was then adjusted to 3 mg/ml using Dulbecco ‘s modified Eagle ‘s medium (DMEM). For the GAG supplemented conditions, either CS, DS (Millipore Sigma), or HA (Millipore Sigma) were dissolved in sterile water to a concentration of 1 mg/ml and then added to collagen during hydrogel preparation, thereby giving a final collagen:GAG ratio of 3:1. This ratio of collagen to GAG constituent was selected based on previous literature for physiological conditions. One example is the ratio of collagen to HA in breast tumors, which range from 2.4:1 – 7.7:1^33,34^. Therefore, the collagen to HA ratio used for this study is in line with levels observed in tumors. After mixing the constituents, the hydrogels were then incubated at 4 °C for 12 minutes to enhance fiber formation.^35^ The incubated collagen-based hydrogels were then injected inside poly(dimethylsiloxane) (PDMS) microchannels that were bonded to glass slides, PDMS wells, or 96 well-plates for their respective assays. Following this, the hydrogels were placed in an incubator (37°C) for 30 minutes. Afterwards, the gels were hydrated with DMEM media and stored at 37 °C for 48 h prior to data acquisition to ensure complete polymerization and equilibrium swelling conditions.

### 2.2. Stiffness Analysis

The mechanical stiffness of the hydrogels was measured after applying a stress-strain relaxation technique adopted from Barocas et al.^36^ Briefly, a mechanical load testing instrument (Electroforce 5500, TA Instrument) was used to indent hydrogels cast in a PDMS well (8mm diameter; 3 mm in height) at a strain rate of 10% per second. The indentation was done in a sequence of four incremental indentations at 10-40% depth, each followed by a 5-minute period of relaxation. From here, the load readings at peaks corresponding to 20-40% strain were used to determine indentation stresses considering indenter geometry. Indentation modulus was calculated after plotting the stresses with their corresponding strain. The data was processed using an in-house MATLAB code. To evaluate the contributions of both the viscous and elastic components of the measured data, the relaxation following each indentation peak was fitted to a viscoelastic model composed of two Maxwell elements (spring in series with a dashpot):

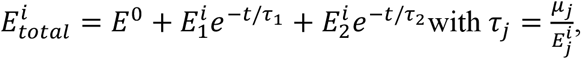

Where 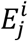 are indentation modulus values, τ_*j*_ are relaxation time constants, μ_j_ are viscosity coefficients, and t is the relaxation time (0-300 seconds for each step). For each indentation step, E_total_ was computed and then fitted to the viscoelastic model.

### 2.3. Microfluidic Hydraulic Permeability Measurement

Hydraulic permeability or Darcy ‘s permeability is a characteristic of the matrix that relates interstitial fluid velocity with an applied pressure gradient.^37^ This measurement was used to characterize the convective transport properties of the collagen hydrogels. The hydraulic permeability was measured using a rectangular microfluidic device (length: 5 mm; width: 500 μm; height: 1 mm) 48 h post hydrogel polymerization as previously described.^24,38^ Briefly, a pressure gradient was created across the channel by fitting a pipet tip filled with DMEM in one port of the microchannel. The hydrostatic pressure gradient generates flow across the hydrogel driven by the height difference between ports. The average velocity of the flow across this channel was then measured by tracking a fluorescent dye (tetramethylrhodamine isothiocyanate−bovine serum albumin (TRITC-BSA, MilliporeSigma), MW= 65.5 kDa) with time-lapse microscopy. Images of the channel were taken on a Nikon TS-100F microscope, equipped with Q-Imaging QIClick camera controlled by NIS elements, every 5 seconds for 5-to-10-minutes. A MATLAB algorithm was then used to estimate the hydraulic permeability from the timelapse videos. Hydraulic permeability (K) was calculated using Darcy ‘s Law:

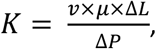

where μ is the viscosity of the fluid (0.001 Pa s), ΔL is the length of the microfluidic channel (5 mm), and ΔP is the hydrostatic pressure gradient across the hydrogel estimated using Pascal ‘s law:

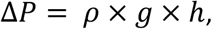

where ρ is the density of the fluid (1000 kg/m^3^), g is the acceleration due to gravity (9.81 m/s^2^), and h is the fluidic height difference between the microfluidic channel ports.

### 2.4. Confocal Reflectance Microscopy for Collagen Fiber Pore Size, Radius, Number, and Solid Fraction Analysis

Confocal reflectance microscopy has been used to observe and characterize the fibers of collagen without the need for fluorescent tagging.^22^ Consequently, subtle changes to collagen microarchitecture due to the addition of GAGs can be imaged and quantified.^24^ Collagen fibers contained within a PDMS microchannel, were imaged using a Nikon A1R live cell imaging microscope via a 40X 1.3 NA oil immersion lens. Exciting the collagen-based hydrogels with a 487 nm laser and then passing the light through a 40/60 beam splitter allowed a detector to collect the light for visualization of the collagen fibers. Approximately eighty to ninety images were collected per stack, each with a z-step of 0.59 μm.

The acquired confocal reflectance stacks were analyzed using a custom MATLAB code developed by our group.^24^ The software determined the fiber radii and pore size of the collagen hydrogels. Fiber radii were measured after skeletonizing the stacks in 3-D using the Skeleton tool of NIH ‘s Image J Software. Total skeleton length was computed to estimate the radius of the fibers. Pore size was estimated using nearest obstacle distance (NOD) method.^39^

To measure collagen solid fraction, the confocal z-stacks were binarized and threshold using ImageJ and processed in an in-house MATLAB code. Briefly, the fiber-occupied space was determined as having a pixel value of “one” and any pore or void space was determined as having a “zero” value. For fiber number, a fiber was determined as any object occupying three pixels with a connecting pixel located at any sixteen possible adjacent face (spaces excluded are placed in the diagonal orientation). This way, the connectivity expanded to the 3-D plane. With the total fiber count and pixel value pertaining to these fibers, solid fraction was determined as the ratio between total pixel volume with fibers and the entire pixel volume (512 × 512 x stacks per image).

### 2.5. Turbidity Assay

Turbidity measurements track collagen aggregate formation by particle light scattering, measuring absorbance as clear hydrogel solutions turn opaque during polymerization.^28^ Collagen-based hydrogels for the turbidity measurements were prepared as described above. Afterwards, 100 μl of hydrogel solution was added to a pre-chilled flat bottom 96-well plate (VWR). The absorbance of the collagen solutions was measured at 400 nm wavelength in an incubated (37 °C) plate reader (Tecan) until the hydrogel polymerized. Separate experiments were conducted to ensure CS, DS, and HA solutions independent of collagen did lead to noise in the turbidity measurements. Turbidity curves are generally sigmoidal and can be divided into three segments: 1) lag phase (or nucleation), 2) growth phase, and 3) plateau phase.^40^ During the lag phase, turbidity is unchanged while during the growth phase turbidity rapidly increases before a plateau is reached as available collagen is depleted.^22^

Turbidity curves were fitted using a sigmoid growth curve

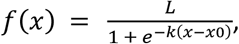

where x_0_ is the sigmoid midpoint, L equals the curve ‘s maximum value, and k is the steepness of the curve. The lag time, inflection time, time to plateau (or plateau time), and the rate of increase in the growth phase (kg), were calculated from the turbidity curves as previously done.^40^ Lag time depicts the start of collagen polymerization and was determined as the point at which the fit to the first derivative curve increased 5% over its initial value. Inflection time defines the halfway point of polymerization and was defined as the time as the midpoint of the sigmoid curve (time of max peak in first derivative curve). Plateau time describes the end of gel formation and equaled the time at which the first derivative curve reached 95% of its maximum value, and kg was described as the linear portion between lag time and plateau time (**Supplementary Figure 1**).

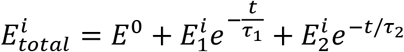

### 2.6. Statistical Analysis

Statistical analysis was conducted using analysis of variance (ANOVA) followed by Tukey ‘s post testing for pairwise comparisons between controls and collagen conditions with CS, DS, and HA. All results shown are presented with error bars representing the standard error of the mean (SEM). Each experiment was run at least with technical duplicates, with 3 sets of biological replicates pooled together for statistical evaluation. A p-value of 0.05 was used as a threshold for statistical significance between results.

## 3. Results

### 3.1. Differential modification of the mechanical properties of collagen-based hydrogels by CS, DS, and HA

First, we determined the bulk mechanical properties of the different hydrogel compositions by measuring the peak indentation modulus using methods previously described by our group and others (**Figure 1A-B**).^24,36^ The indentation modulus was significantly greater by 2.3-fold for the collagen + HA condition (5.05 kPa) versus the collagen only condition (2.16 kPa) (**Figure 1C**). In contrast, the indentation modulus for collagen + CS (2.35 kPa), collagen + DS (2.35 kPa), and collagen only were not statistically different. While both CS, DS, and HA have been shown to resist compressive loading, our results suggest that HA but not the CS and DS exerts a dominant effect in the resistance to compression when integrated with fibrillar collagen-based hydrogels. Further analysis into the relaxation conditions of the hydrogels via the modeling with the Maxwell elements (**Supplemental Figure 2**), showed that while the GAGs impart different bulk mechanical properties, with HA primarily being the sole mediator of stiffness as conferred by indentation, the materials are all relaxing within the same time scale (**Supplemental Figure 2**). Recently it was reported that the relaxation of hydrogel materials is pivotal in mediating cell motility and adhesion.^41^ Notably, collagen + DS and collagen + HA have a higher relaxation load under strain when compared to other conditions. For nonfibrillar matrix components, this aspect is normally attributed to resistance to volume change when subjected to compression.^36^

**Figure 1:**
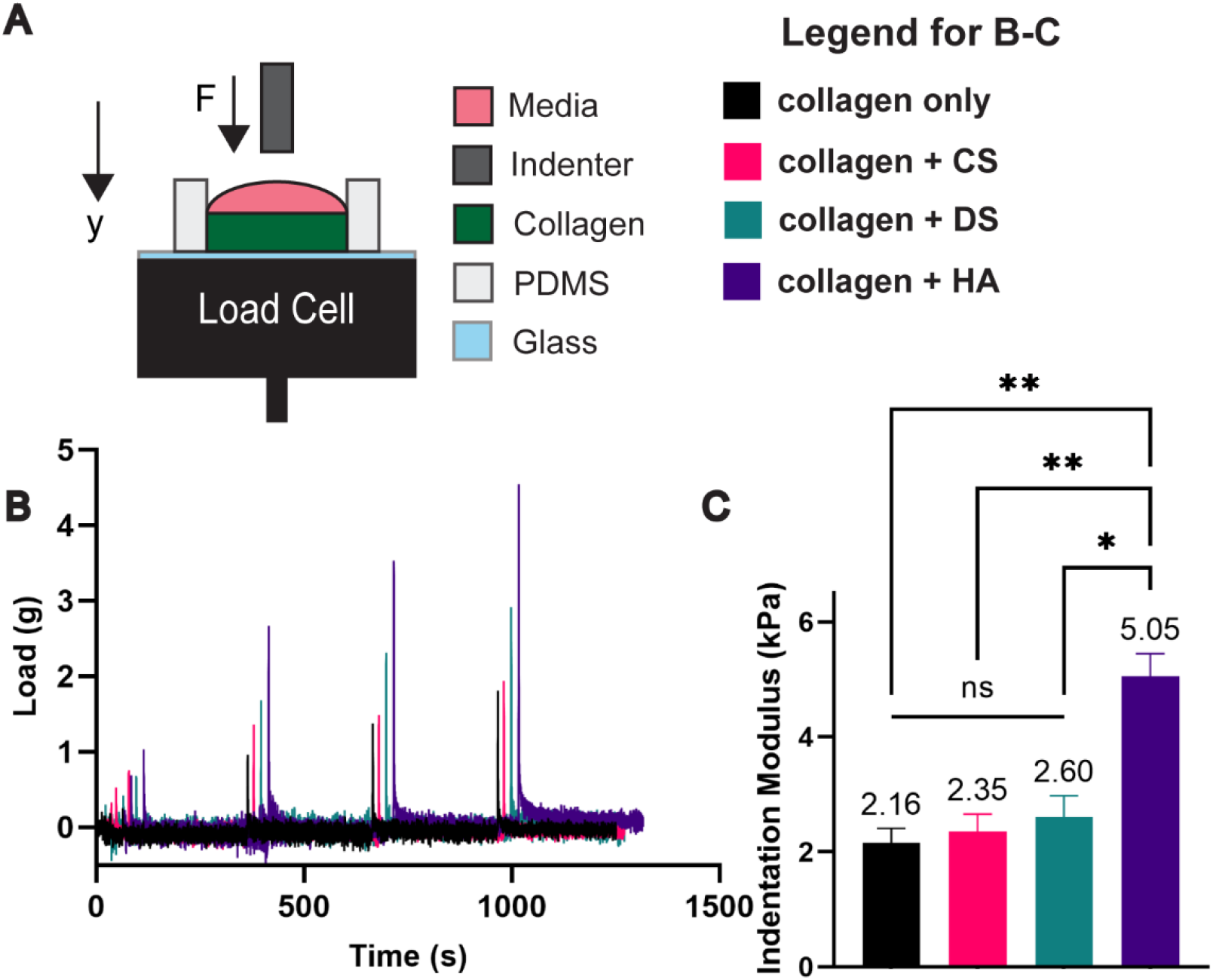
Differential effects of CS, DS, and HA in altering hydrogel stiffness. A) Collagen solutions are cast on a PDMS well and subjected to stresses using indentation. B) Sample load readings of tested conditions. C) Only HA laden hydrogels led to increases in stiffness. Error bars denote SEM; NS is non-significance, *<0.05, **< 0.01.

### 3.2. CS, DS, and HA do not significantly alter the hydraulic permeability of collagen-based hydrogels

Next, we evaluated the convective transport properties of the hydrogels as these are dependent on ECM composition.^24^ Mass transport properties (i.e., convection and diffusion) influence cell behavior by controlling physiologically-important parameters such as shear stresses, chemical gradients, nutrient availability, and drug delivery.^37,42^ Following a protocol developed by our group, a microfluidic device was used to evaluate the effect of CS, DS, or HA on hydraulic permeability, which is a coefficient that describes the convective transport properties of semiporous medium such as the ECM.^43^ The addition of either CS, DS, or HA to collagen hydrogels showed no significant effects on hydraulic permeability when compared to collagen alone (**Figure 2**). Previous work from our lab attributed this outcome to the competing efforts between increases in fiber radius and increases in pore size when adding HA to collagen matrices, which nullified any potential changes in permeability due to increasing GAG content.^24^ This outcome was supported by a past study that demonstrated that a GAG rich hydrogel led to decreased transport when compared to a collagen-GAG solution. ^44^ In a collagen-GAG mixture, GAGs such as HA and CS tend to associate around the collagen fibers and become excluded from the interstitial space which may explain the nonsignificant change in permeability.

**Figure 2:**
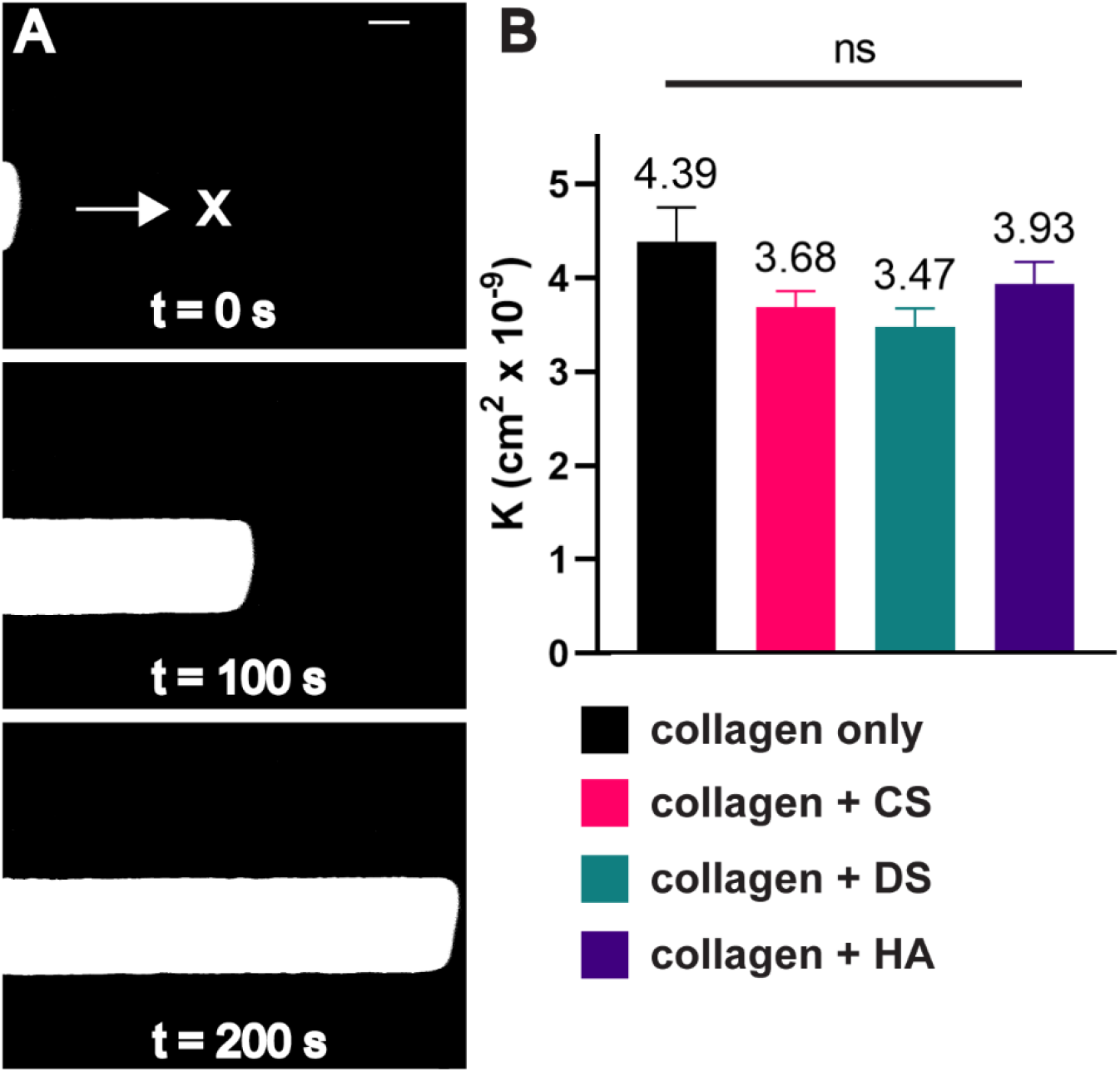
Hydraulic permeability measurements of GAG modified collagen matrices. A) Time-lapse microscopy is used to track a TRITC dye in a microfluidic channel to obtain the velocity of the flow. B) The addition of HA, CS, or DS fails to decrease hydraulic permeability (K). Scale bar = 200 μm. Error bars denote SEM. NS is non-significant ANOVA for all conditions tested.

### 3.3. Differential effects of CS, DS, and HA on collagen matrix microstructural properties and fiber content

Next, we used confocal reflectance microscopy (**Figure 3A**) to quantify how modifications to collagen polymerization kinetics led to differences in the collagen matrix microarchitecture parameters of fiber radius, pore size, solid fraction, and total fibers (**Figure 3B-E**). Compared to collagen only, collagen + DS and collagen + HA significantly increased fiber radius by ∼22% (DS) and 12% (HA) (**Figure 3B**). Similarly, collagen + DS and collagen + HA increased matrix pore size by 24% and 15% respectively when compared to collagen only (**Figure 2C**). Interestingly, and in contrast to collagen + DS and collagen + HA, collagen + CS had no significant effect in fiber radius (**Figure 3B**) and significantly decreased pore size by 8.0% (**Figure 3C**) when compared to collagen only. In **Figure 3D** we plot the solid fraction. The volume fraction of solids in fibrillar collagen-based matrices are a property of fiber radius (**Figure 3B**), fiber length, and the number of fibers. When compared to the collagen only condition, collagen + CS and collagen + DS had no significant effect on the solid fraction. In contrast, collagen + HA had a significant decrease of ∼ 20% in solid fraction compared to collagen only **(Figure 3D**). In **Figure 3E**, we plot the total fiber count per z-stack as a measure of the number of fibers. The collagen + CS matrix condition significantly increased total fibers by ∼ 14% when compared to the collagen only condition **(Figure 3E)**. In contrast, when compared to collagen only, collagen + DS and collage + HA significantly decreased total fibers by 35% and 17% respectively.

**Figure 3:**
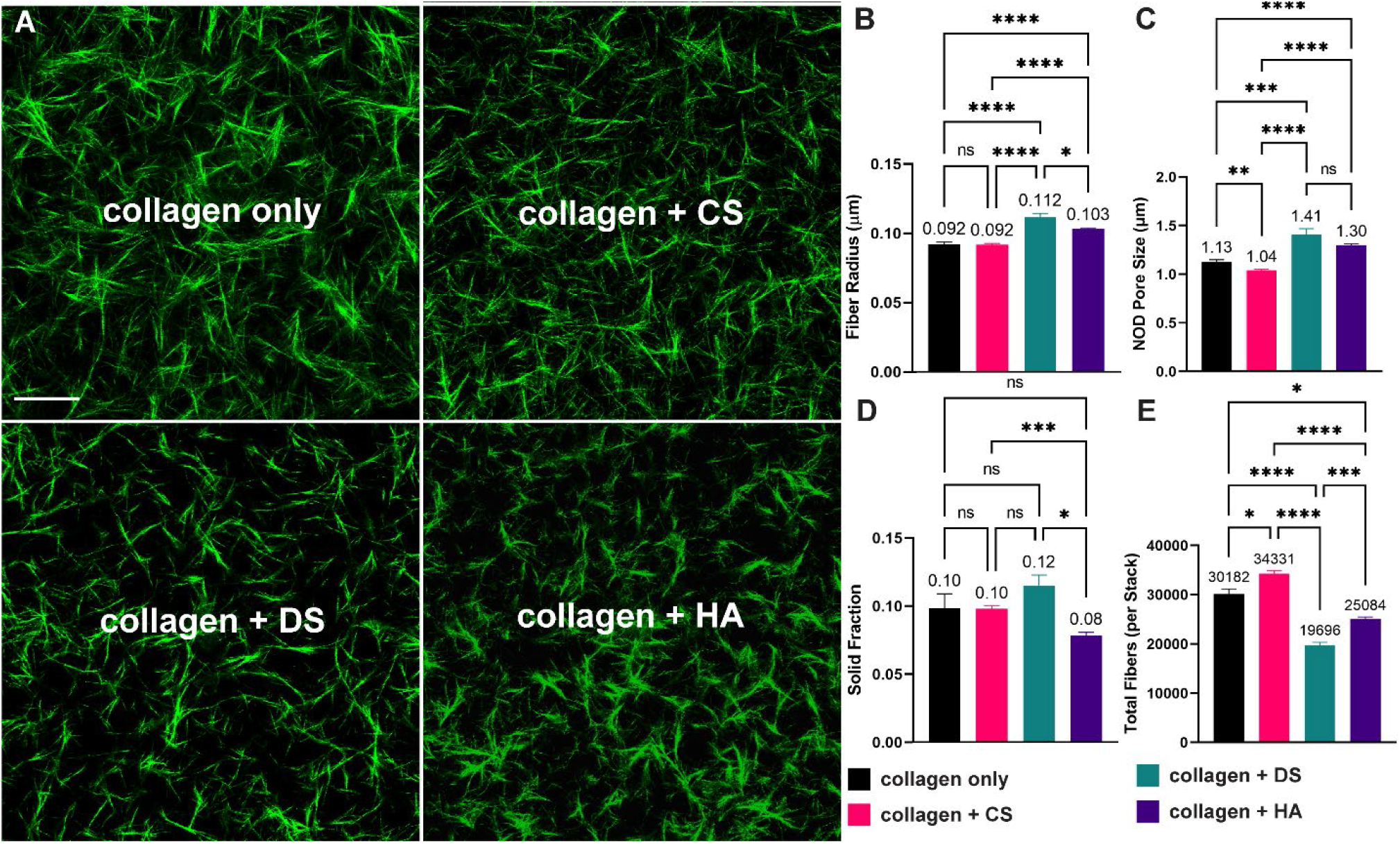
Microstructural analysis of collagen-based hydrogels. HA and DS increase both matrix pore size and fiber radius, while CS leads to decrease in fiber pore size. A) Representative images of collagen fibers. B-C) Addition of DS and HA leads to increases in pore size and radius. D) HA decreased the solid fraction following processing of confocal images. E) DS and HA decreased the number of collagen fibers, while CS led to increased amounts of fibers when compared to collagen only. Scale bar = 50 μm. Error lines represent the SEM. Statistical significance denoted as NS is non-significance, * <0.05, **<0.01, *** <0.001, **** < 0.0001.

### 3.4. CS, DS, and HA differentially influence the rate of collagen polymerization

Next, we used turbidity measurements to evaluate how GAG supplementation affects the polymerization dynamics of collagen matrices. (**Figure 4A**). Interestingly, while the maximum optical densities (ODs) during the plateau phase for the collagen + CS, and collagen + DS conditions were within +/-2.2%, the maximum OD for the collagen-only condition, the OD for the collagen + HA condition was ∼ 13% lower than the collagen-only condition (**Figure 4A**). This outcome suggests that the collagen + HA condition has less total polymerized collagen fiber content compared to other experimental conditions. This result also agrees with the solid fraction (**Figure 3D**) and total fiber number (**Figure 3E**) results. Further analysis of the polymerization kinetics showed that collagen + CS did not change the lag time statistically compared to collagen only (**Figure 4B**) but significantly decreased the inflection time (495 s versus 577 s, **Figure 4C**) and the plateau time (759 s versus 907 s, **Figure 4D**). In contrast, when compared to collagen-only, collagen + DS significantly decreased the lag time (296 s versus 348 s) and the inflection time (529 s versus 577 s) but did not change the plateau time (**Figure 4B-D**). Our results with CS and DS are in accordance with previous studies, which have shown that CS modify collagen fiber microstructural properties.^45,46^

**Figure 4:**
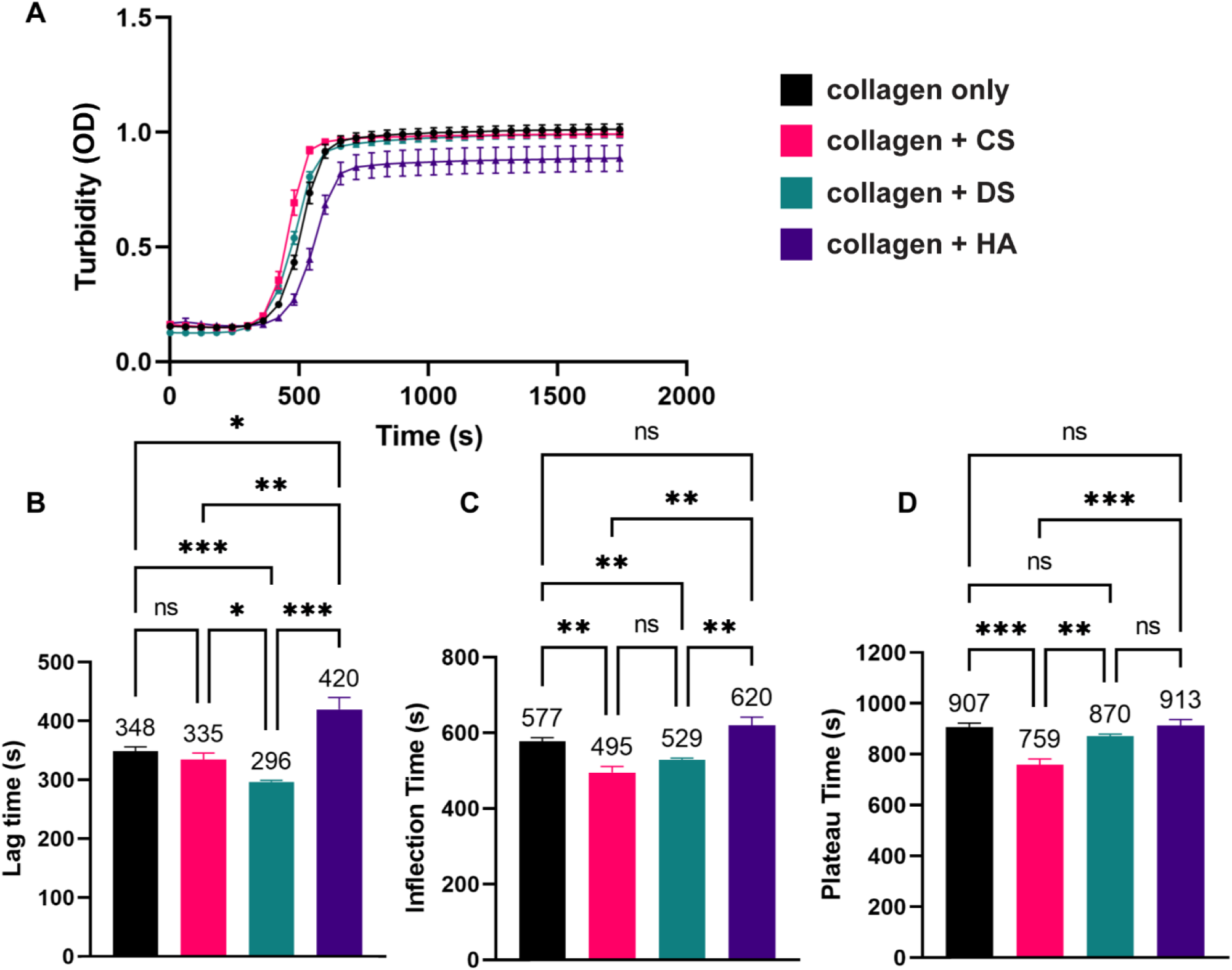
Turbidity measurements of collagen gel solutions. A) Addition of GAGs differentially affects the rate of collagen fibril polymerization in terms of B) Lag time, C) Inflection time, and D) Plateau time. While CS and DS enhance the rate of collagen fibril formation, HA hinders the rate of polymerization. Error lines represent the SEM (n > 9 per condition). Statistical significance denoted as NS is non-significance, *<0.05, ** <0.01, *** < 0.001.

Compared to the collagen-only condition, collagen + HA significantly increased the lag time (420 s versus 348 s) but did not change the inflection and plateau times (**Figure 4B-D**). The observation that HA delays the lag phase of collagen polymerization is in agreement with previous results, which described that HA sterically impedes the aggregation of collagen molecules during self-assembly.^6^ Interestingly, collagen + CS resulted in a significant increase in the growth phase (kg), compared to the collagen-only, collagen + DS, and collagen + HA conditions (**Supplemental Figure 1**). These outcomes suggest that CS but not DS and HA promotes more rapid collagen fibrillogenesis compared to collagen-only conditions.^47^

## 4. Discussion

In the design and utilization of hydrogels for cell culture and tissue engineering applications, it is essential to characterize the properties that influence the compositional, mechanical, and structural cues to cells. In this study, we quantified the biophysical properties of collagen-based hydrogel systems using an integrated approach that enabled independent measurements of key mechanical, transport, and microarchitectural characteristics. Our studies focused on hydrogels comprised of physiological collagen:GAG ratios. For all experiments, we used a fixed collagen concentration (3 mg/ml) and compared outcomes of the addition of three different GAG types (CS, DS, and HA) at 1 mg/ml. Our results provide a uniquely detailed understanding of how the addition of different GAG types differentially alter collagen polymerization and turbidity kinetics that translate to distinct physical and structural properties of the ECM. These results will aid researchers who aim to gain a better understanding of the combined biological effects of multiple ECM parameters, such as composition and biophysical properties, on cell behavior and function.

Turbidity measurements enable temporal distinctions between the collagen and GAG conditions during hydrogel polymerization. The optical density of the absorbance measurements that correspond to the plateau in the sigmoid turbidity curves relates to the total collagen content present in fibril form.^48^ Our turbidity measurements are supported by the microstructural features and the collagen solid fraction present in the matrices studied. For instance, the inflection and plateau times for collagen + CS from the turbidity curves were significantly decreased compared to collagen-only. This result suggests that the addition of CS accelerated total collagen polymerization. More rapid collagen fibrillogenesis and fiber count due to CS addition is further supported by the rate of increase during the growth phase (slope between plateau and lag time) of the sigmoid turbidity curves. Examining the microstructural features, the significant increased number of total fibers for collagen + CS compared to collagen only also suggests that CS enhances collagen fibrillogenesis. Thus, for collagen + CS, we observed increased fiber content, with reduced pore size and no change in fiber radius compared to collagen only, which suggests that the addition of CS modifies the length of collagen fibers or the total number of pores.

For the collagen + DS condition, we observed in the turbidity measurements decreased lag and inflection time when compared to collagen only. Shifts to the left of the sigmoid turbidity curve has been reported to be associated with accelerated fibrillogenesis^25^, which corresponds with fibril thickening.^40^ This outcome is supported by the microarchitecture analysis for collagen + DS, where we observed increased radius, and collagen solid fraction.

In the collagen + HA condition, although the addition of HA delays the start of collagen polymerization, the ratio between plateau time and lag time is comparable to CS, which has the most rapid polymerization of the conditions tested. These results suggest that while HA can sterically impede the start of collagen aggregation and bundling, once hydrogel polymerization initiates, the process proceeds to accelerate. It is important to note that while HA increases fiber radius – denoting enhanced polymerization – it also decreases the total number of fibers, which can explain why less time is needed for the hydrogel to fully polymerize. Recently, Martin *et al*., described the effects of the simple monosaccharides D-glucuronic acid, N-acetyl-D-glucosamine, and D-galactose in modifying collagen assembly.^49^ This study observed that either the presence of amide groups or the charged nature of carboxylate groups associated with the sugars (D-glucuronic acid) interact with water molecules on the surface of collagen monomers, thereby increasing the entropy change necessary for spontaneous collagen assembly. While this study did not investigate complete GAG chains, the results provide a plausible explanation for the delay in collagen fibrillogenesis in the presence of HA. Since HA is composed of repeated N-acetyl-D-glucosamine and D-glucuronic acid chains, the molecule is having a similar effect on collagen self-assembly compared to the isolated sugars. Future work can focus on the entropy changes caused by the addition of sulfated GAG chains like CS or DS that impart a more negative charge,^18,50,51^ to fully understand the influence of GAGs on the kinetics of collagen assembly.

Structurally, CS, DS, and HA are similar anionic polysaccharides. CS is known to have a higher water retaining capacity than HA, leading to ECM with significant swelling that contributes to load-bearing behavior of CS rich tissue such as cartilage.^50,52^ However, of the conditions tested, only collagen + HA hydrogels exhibited increased indentation modulus. Interestingly, this result was not contingent on the solid collagen fiber fraction as in the HA condition, both collagen solid fraction and total fiber count decreased. Our data suggests that as a component of collagen + HA compositive hydrogel, HA promotes resistance to compression presumably due to sustained swelling while being indented. For CS and DS, the collagen solid fraction increased when compared to HA conditions; however, this fails to increase the modulus. With CS, DS, and HA causing swelling in collagen matrices, it was unexpected that only the addition of HA led to significant increases of the indentation modulus with no discernable link between fiber radius, total fiber, and solid fraction. Yet, it is notable that in a previous study, it was shown that enzymatic degradation of CS with chondroitinase did not change the compressive modulus of a collagen-CS gel.^53^ In contrast, cleaving HA with hyaluronidase significantly reduced the compressive modulus of a collagen-HA gel to the level of a collagen only.^24,31^ Collectively, these results suggest additional factors may influence the mechanical behavior of collagen-GAG composite hydrogels that are either experimental parameters (e.g., strain rate or level of confinement) or properties of the material tested (e.g., degree of cross-linking of the biopolymer networks).^50^

Hydraulic permeability correlates to the drag imparted by the solid compartment of the ECM, with both fibrous and nonfibrillar components contributing to hydraulic resistance.^37^ Carman-Kozeny and Happel describe hydraulic permeability further in terms of the geometry of the channel, the overall porosity, and the hydraulic radius of the ECM pores, all of which play a role in defining permissiveness to fluid flow through semiporous matrices.^37,54,55^ For HA, both the solid fraction and number of fibers decrease, which combined with the pore size increase would be counteracted by the increase in fiber radius. Similarly, DS leads to increases in pore size and radius, with decreasing number of fibers, but the total solid fraction is increased suggesting that there could be excess drag associated with the increased collagen fiber content (either through decreased amounts of pores or increased fiber length). Finally, CS decreases pore size but has no change in fiber radius with comparable solid fraction to the collagen only condition. Moreover, as stated previously, in a developing biopolymer network, GAGs may associate around individual collagen fibers instead of occupying the interstitial space.^44^ Therefore, while it is surprising that the addition of the GAGs does not affect hydraulic permeability, all GAG-laden conditions changed at least one fiber property tied to hydraulic permeability.

Despite observing no significant changes in hydraulic permeability of the hydrogel conditions tested, we believe that these findings will still be very useful for experimental studies that focus on the effects of ECM GAGs on cellular responses, especially when combined with other biophysical measurements such as microstructural analysis. For instance, increases in the mean pore size of collagen-only matrices may facilitate cancer cell invasion.^26^ In addition, collagen fibers that define the pore size are sufficiently deformable to support angiogenic sprouting of endothelial cells.^56,57^ Thus, our findings suggest that the significant increases in collagen matrix pore size due to HA or DS addition may facilitate the initial extension and sustained elongation of multicellular processes into matrix void spaces during cancer invasion or endothelial sprouting. Indeed, we previously reported that HA addition to collagen ECM promoted angiogenesis in a microfluidic-based model but in the presence of convective interstitial flow only.^31^ Thus, modifications in the matrix pore size due to HA addition resulted in changes in interstitial flow induced-shear stress on endothelial cells, which promoted angiogenic sprouting. The utility of the hydraulic permeability measurements is that we can conclude that these changes observed in endothelial sprouting were not dependent on changes in nutrient limitation, since hydraulic permeability was unchanged with the addition of HA. Like HA, CS and DS have pathophysiological relevance.^58^ Thus, we believe that the findings from this study will be instrumental for gaining a more complete understanding of the contributions of the myriad of instructional cues to cells encoded by collagen ECM enriched with GAGs, especially in the presence of physiologically-relevant interstitial flow conditions.

We anticipate that the integrated characterization scheme for hydrogels presented here will have broad application for cell culture and tissue engineering studies, especially ones using collagen-based matrices. For example, we can readily characterize the biophysical, solid fraction, and turbidity properties of collagen-based matrices whose stiffness is modified independent of collagen concentration using methods such as but not limited to enzymatic or chemical cross-linking,^59^ non-enzymatic glycosylation (or glycation),^60^ or non-enzymatic cross-linking of collagen/alginate hybrid gels.^61^ Moreover, we can also study modified collagen-GAG composite materials, such as ones that utilize commercially available methacrylated collagen and HA for covalent crosslinking used to slow down hydrogel degradation and enhance mechanical tunability.^29,62^ However, it is known that HA methacrylation not only changes hydrogel bioactivity but alters collagen microstructure.^32^ To our knowledge, such collagen-based matrices (i.e., independently stiffness tuned and methacrylated collagen-HA) have not been widely deployed in microfluidic systems that combine controlled perfusion and interstitial flow with monitoring and manipulating of cellular environments in real time.^43^ Thus, the approaches described here would help position researchers to investigate the interrelationship between mechanical and transport factors mediated by interstitial flow with ECM-derived biochemical and biophysical cues in orchestrating biological function.

## 5. Conclusions

In this study, we employed an experimental framework that described the effects of the GAG molecules CS, DS, and HA in differentially modifying the properties of collagen-based matrices. Here, we find that the GAGs interact with the collagen fibers during polymerization that manifest in distinct ECM biophysical properties. Thus, the addition of different GAGs not only distinctly alters the bioactivity of collagen-based matrices, but they also change the mechanical and structural properties to define the initial conditions for reciprocal interactions between the matrix and cells. Complementary investigation of the structural, mechanical, and turbidity properties of developing biopolymer networks, as has been performed in this study, opens new avenues for precise understanding of the complex gelation processes of collagen-GAG composite hydrogels. Such investigation will enable enhanced design and utilization of biomaterials for cell culture and tissue engineering applications. Moreover, studies detailing the interrelation between compositional, microarchitectural, and mechanical properties will allow for a fuller understanding of how cells and biophysical environments of the ECM function an integrated system to influence cell and tissue behavior and response.

## Acknowledgements

The authors acknowledge support from an NSF CAREER award (CBET-1752106), the Mark Foundation for Cancer Research (18-024-ASP), the National Heart Lung Blood Institute (R01HL141941), and The Ohio State University Materials Research Seed Grant Program, funded by the Center for Emergent Materials, an NSF-MRSEC, grant DMR-1420451, the Center for Exploration of Novel Complex Materials, and the Institute for Materials Research. Two of the authors (P.E.B. and M.M.M.) gratefully acknowledge funding from the OSU (Ohio State University) Pelotonia fellowship program. One of the authors (M.C-M.) thanks the support from an OSU Graduate Enrichment Fellowship, a Discovery Scholars Fellowship, and an NHLBI Graduate Diversity Supplement. J.J.A is supported by funding from the GEM Fellowship, Ohio State University Discovery Fellowship, and the Gates Millennium Scholarship. S.S.A is funded through the Ohio State University Fellowship and the Ohio State Distinguished University Fellowship. We thank Jacob Holter for critical review of this manuscript.

