## Supplementary FIgures for "Chondroitin sulfate, dermatan sulfate, and hyaluronic acid differentially modify the biophysical properties of collagen-based hydrogels"

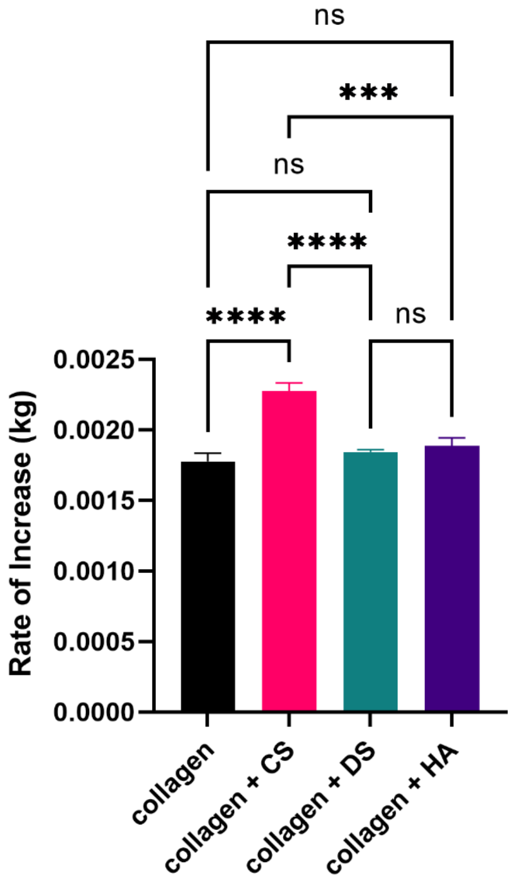


Supplementary Figure 1: Rate of Increase (kg). CS increases the linear slope between lag time and plateau time (~steepness of the sigmoid curve). Error bars denote SEM in the graph. NS is non-significance, *** < 0.001; **** < 0.0001.

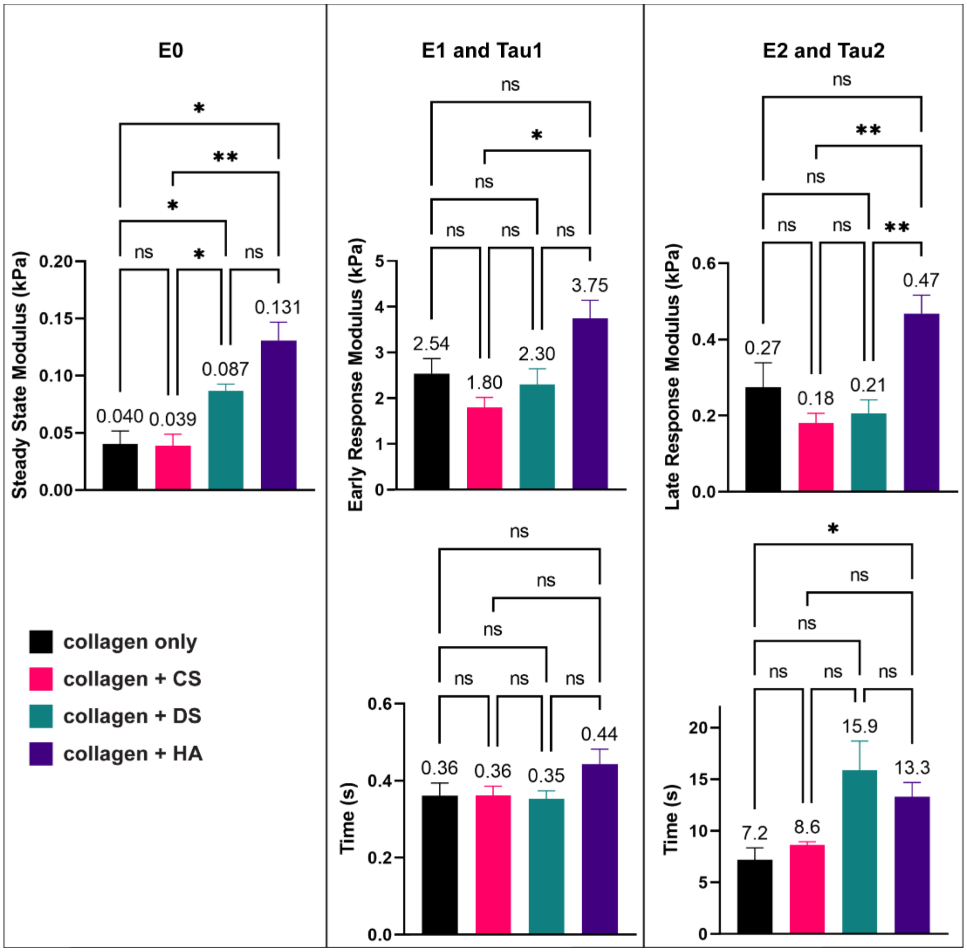


Supplementary Figure 2: Two-Relaxation Time Solid Linear Model ($E_i$ = Elastic Moduli; $Tau_j$ = Relaxation Time Constants. Results for the two-time relaxation solid linear model show minimal changes in the viscoelastic behavior of the different hydrogels for 40% strain. Error bars denote SEM. NS is non-significance, **< 0.01, * < 0.05.

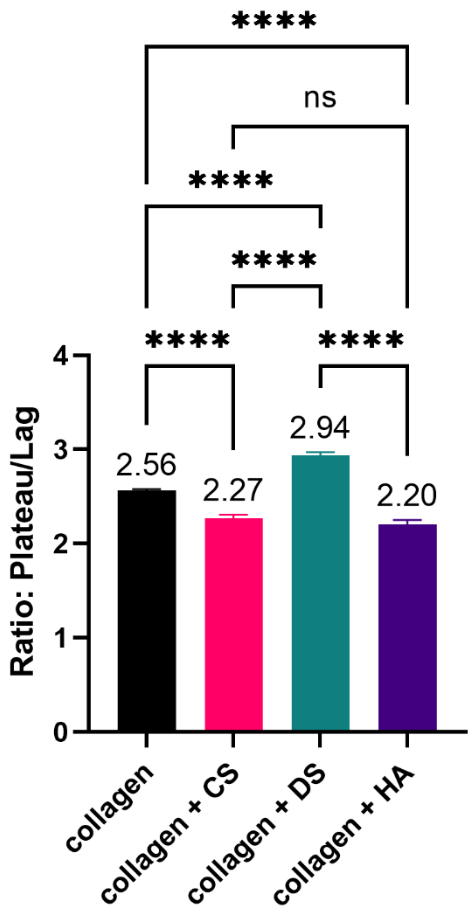


Supplementary Figure 3: Ratio between Plateau Time and Lag Time from Sigmoidal Turbidity curve. CS and HA decrease the time from lag to plateau when compared to collagen + DS and collagen only conditions. Error bars represent the standard error of the mean (SEM). NS is non-significance, p of **** < 0.0001.

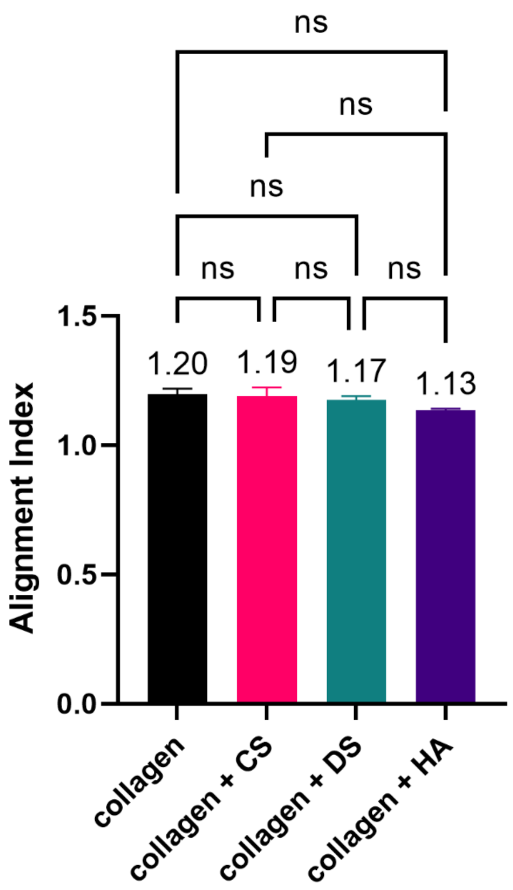


Supplementary Figure 4: Alignment Index. Alignment of collagen fibers quantified showed no difference in the alignment index with the addition of the GAGs. Error bars denote standard error of the mean. NS is non-significance.
